# Using UAV-based temporal spectral indices to dissect changes in the stay green trait in wheat

**DOI:** 10.1101/2023.07.14.549080

**Authors:** Rui Yu, Xiaofeng Cao, Jia Liu, Ruiqi Nie, Chuanliang Zhang, Meng Yuan, Yanchuan Huang, Xinzhe Liu, Weijun Zheng, Changfa Wang, Tingting Wu, Baofeng Su, Zhensheng Kang, Qingdong Zeng, Dejun Han, Jianhui Wu

## Abstract

Stay green (SG) in wheat, a beneficial trait for increasing yield and stress resistance, needs to be supported by analysis of the underlying genetic basis. Spectral reflectance indices (SIs) provide non-destructive tools to evaluate crop temporal senescence. However, few SI-based SG quantification pipelines for analyzing diverse wheat panels in the field are available. Here, we first applied SIs to monitor the senescence dynamics of 565 diverse wheat accessions from anthesis to maturation stages during two field seasons. Based on over 12,000 SIs data set, four SIs (NDVI, GNDVI, NDRE and OSAVI) were selected to develop relative stay green scores (RSGS) and the senescence of wheat populations occurs mainly at four developmental stages stage 1 (S1) to S4, accounting for the final SG indicators. A RSGS-based genome-wide association study identified 47 high-confidence quantitative trait loci (QTL) harboring 3,079 SNPs significantly associated with RSGS and 1,085 corresponding candidate genes in the two seasons; 15 QTL overlapped or were adjacent to known SG-related QTL or genes and the remaining QTL were novel. Finally, we selected three superior candidate genes (*TraesCS6B03G0356400*, *TraesCS2B03G1299500*, and *TraesCS2A03G1081100*) as examples by transcriptomes, gene annotation, and gene-based association analysis for further analysis and found that utilization of superior SG-related variation in China gradually increased following the Green Revolution. The study provides a useful reference for further SG-related gene discovery of favorable variations in diverse wheat panels.

## 1. Introduction

Wheat varieties with high and stable yield are required to meet the demand of a growing world population and a changing climate (Xiao et al., 2022). Senescence in wheat is a highly complex developmental process that progresses from flowering until harvest ripeness and is closely related to the final yield (Sultana et al., 2021, Kamal et al., 2019). Senescence reduces yield when it occurs prematurely due to genetic and adverse environmental factors, such as disease, heat, drought, and low nitrogen conditions (Sultana et al., 2021, Kamal et al., 2019, Gregersen et al., 2013, Kumar et al., 2021, Distelfeld et al., 2014). Stay green (SG) is a dynamic phenotype used to describe the maintenance of post-anthesis canopy vigor that delays senescence and prolongs photosynthetic activity during grain filling (Munaiz et al., 2020, Thomas & Ougham, 2014). SG in wheat is positively correlated with the yield and is associated with stress tolerance (Kumar et al., 2021, Kipp et al., 2014, Thomas & Ougham, 2014). SG is a complex quantitative trait and an understanding of the genetic basis will provide selective markers for use in breeding.

In recent years, SG-related QTL or genes, such as *Stg1* to *Stg4*, *Brnye1*, *CsSGR*, *NAC7* and *OsSGR*, have been detected by genome-wide association studies (GWAS) or quantitative trait loci (QTL) mapping in sorghum, cabbage, cucumber, maize and rice (Crasta et al., 1999, Borrell et al., 2022, Wang et al., 2018, Wang et al., 2019, Zhang et al., 2019, Shin et al., 2020). These genes are involved in phytohormone, transcriptional factor, and multi-enzyme functions and provide a preliminary understanding of the forward genetics involved in SG-related molecular mechanisms. In common wheat, combining recombinant inbred lines (RILs), doubled haploid (DH) populations, multi-reference nested association mapping (MR-NAM) populations and QTL mapping have identified some SG-related QTL (Ren et al., 2022, Shi et al., 2017, Christopher et al., 2021), but no further genetic exploration has been conducted. In tetraploid wheat, the senescence-promoting factor *NAM-B1* was explored by QTL mapping, reduction in RNA levels of the multiple *NAM* homologs delayed senescence (Uauy et al., 2006). Moreover, some transcription factors were identified that bind to plant hormones to stay green through mutant identification and gene differential expression in Arabidopsis, such as *NAC* and *WRKY* (Guo et al., 2021). In recent study, *TaARF15-A1* was identified as a negative regulator of senescence in wheat, it suppressed the expression of *TaNAM-1* via protein–protein interaction and competition with *TaMYC2* for binding to its promoter to regulate senescence (Li et al., 2023). However, there are further genetic factors regulating SG that need to be identified (Kusaba et al., 2013). Recent advances in genome sequencing provide new opportunities to understand the genomic structure underlying SG in wheat (Walkowiak et al., 2020). Successful GWAS or QTL mapping of functional genes requires high-quality genomic and phenotypic data. Although access to high quality genomics data has become much easier, acquisition of appropriate and accurate SG phenotype data is lagging far behind (Song et al., 2021).

Unmanned aerial vehicle (UAV)-based temporal spectral indices have replaced traditional phenotypic acquisition methods and are widely used in evaluating SG traits (Hassan et al., 2021, Liedtke et al., 2020). They allow development of supporting phenotypic pipelines to quantify SG phenotype(Hassan et al., 2021, Liedtke et al., 2020, Christopher et al., 2021). However, these pipelines are suitable only for small groups of genotypes with relatively consistent phenology (such as heading or flowering date). Florescence is generally inconsistent among individuals in diverse wheat panels due to differences in flowering time especially when comparisons are made using a single stage of development or absolute senescence responses of individual accessions. Few studies have considered this issue and have quantified SG in diverse wheat panels using temporal SIs for SG-related gene detection. An effective pipeline encompassing multi-stage SG trait quantification needs further exploration to address this issue.

The present work aimed to develop a new phenotyping pipeline to identify genetic variation in SG in diverse wheat panels under field conditions. We identified wheat SG-related genes that were selected during domestication and in modern breeding. We also identified favorable alleles that were more intensively selected in different wheat regions.

## 2. Materials and Methods

### 2.1 Plant materials

A panel of 565 diverse wheat accessions including Chinese landraces (CL), introduced (foreign) modern cultivars (IMC), modern Chinese cultivars (MCC), and breeding lines from our previously collected ∼5,000 accessions were used for phenotyping (Wu et al., 2021). Detailed descriptions of the accessions are provided in Table S1. This panel was subjected to genotyping with the 660 K SNP array (Beijing CapitalBio Technology Company, http://www.capitalbiotech.com) and used for genome-wide association mapping. Further details on the quality control of genotype data in this panel were reported in Wu et al. (Wu et al., 2021). A second panel of 584 available genome re-sequences based on published data at website http://wheatomics.sdau.edu.cn/ was mainly used for candidate gene-based association analysis (Ma et al., 2021). A pair of wheat cultivars Shaannong 235 and Shannong 15 with the contrary phenotype (stay green/non-stay green) were used for RNA-seq analysis.

### 2.2 Field experimentation

The panel of 565 diverse wheat accessions was planting at Cao Xinzhuang Experimental Farm at Yangling (34.31°N, 108.10°E) in Shaanxi province of China during the two cropping seasons (2020-2022) (Fig. 1). The experiments were arranged in an augmented design with 16 blocks and five controls repeated in each block by ACBD-R (Burgueño et al., 2018). Each plot was 6 m^2^ (6 × 5 m rows with 20 cm spacing and plant density of 2.7 million/hectare. Reflectance measurements were made from 50% anthesis until harvest ripeness. Daily air temperature and rainfall data during each test period were obtained from the Yangling National General Weather Station (Fig. S1).

**Fig. 1.**
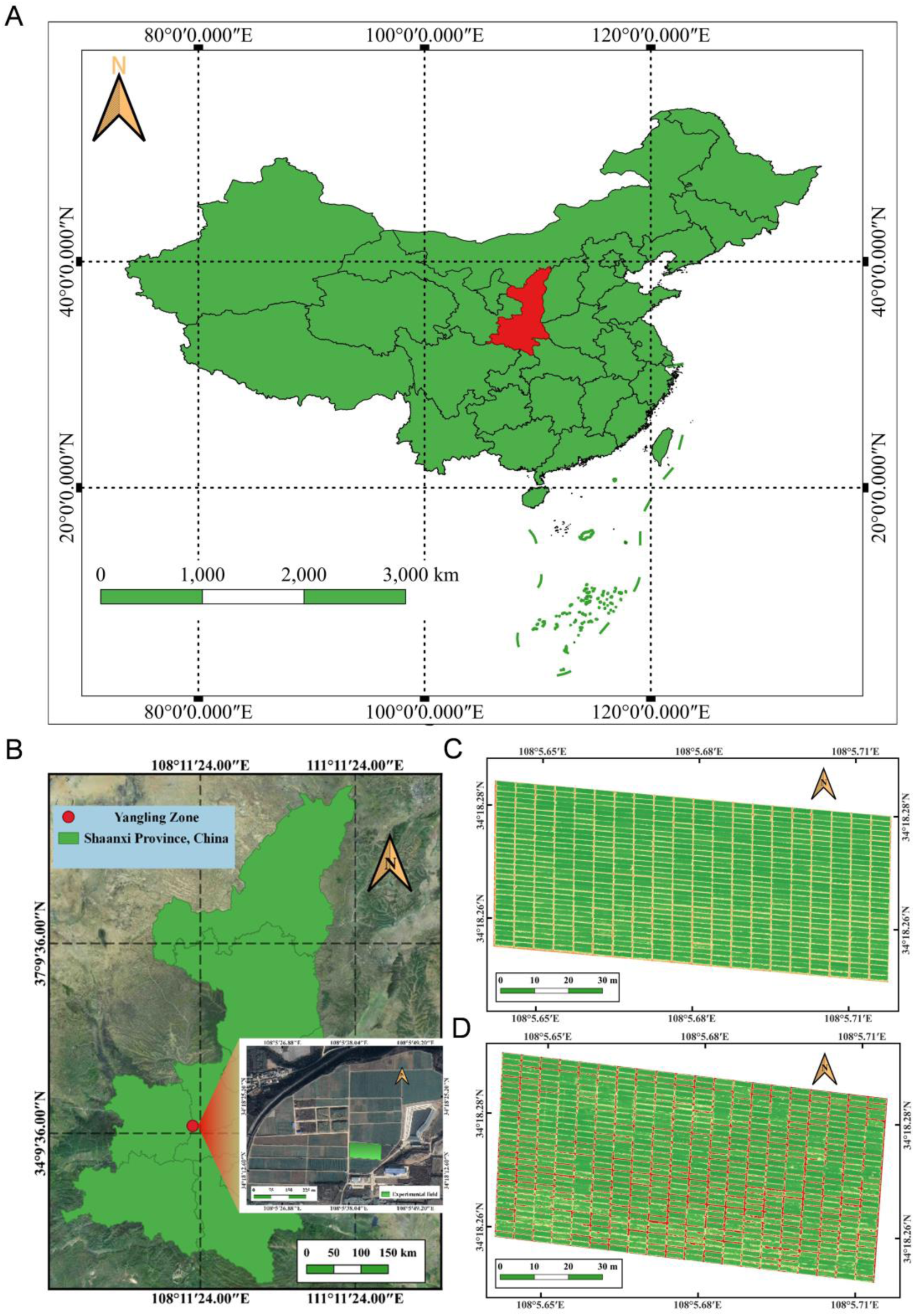
(A) Map of China. (B) Yangling Zone, Shaanxi Province. Location of Cao Xinzhuang Experimental Farm and the experimental field (lower right corner). Orthophoto map of wheat population at anthesis stage in 2020-2021 season (C) and 2021-2022 season (D).

### 2.3 Acquisition of temporal multispectral images

A MicaSense Altum camera (Micasense, USA) was used to collect data. The camera was mounted on a DJI Matrice 200 UAV (DJ-Innovations Technology, China) in 2020-2021 and a DJI Inspire 2 UAV (DJ-Innovations Technology) in 2021-2022. The camera encompassed five narrow spectral bands and the specific information for each band is shown in Table 1. The spectral bands have a 2,064 × 1,544 (pixel) spatial resolution and a ground sample distance (GSD) of 5.28 cm per pixel at a height of 120 m. A Downwelling Light Sensor 2 was used to correct for ambient light changes during flight. A Global Position System (GPS) sensor was used to record the geographic location information of images. A calibrated reflectance panel (CRP) was photographed before each flight to calibrate band reflectance. The CRP has an absolute reflectance, making it possible to compare images from different flights. Temporal images were acquired at 2 - 4 days intervals in full sunshine at mid-day from anthesis to maturity. UAV spectral data were obtained in 10 and 9 flights in 2020-2021 and 2021-2022, respectively. The flight height to the wheat canopy was 26-27 m and the forward and side overlap rates were 85-90%.

**Table 1.**
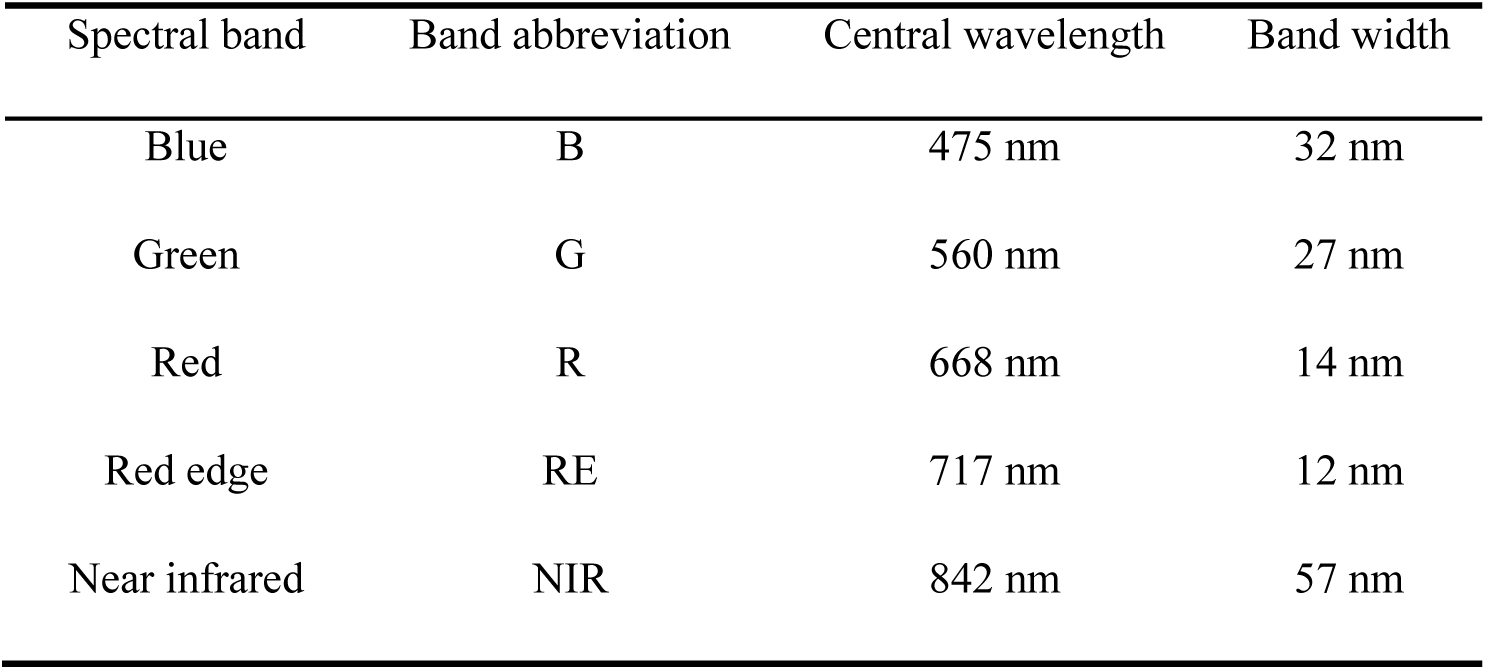
Spectral specifications of the MicaSense Altum camera.

| Spectral band | Band abbreviation | Central wavelength | Band width |
| --- | --- | --- | --- |
| Blue | B | 475 nm | 32 nm |
| Green | G | 560 nm | 27 nm |
| Red | R | 668 nm | 14 nm |
| Red edge | RE | 717 nm | 12 nm |
| Near infrared | NIR | 842 nm | 57 nm |

### 2.4 Image processing and index extraction

The captured images from each flight were processed to generate reflectivity orthophoto maps for each band using Pix4D Mapper (Pix4D, Switzerland). The geographical deviations of orthophoto maps for different flights were calibrated based on ground control points in the Quantum GIS (QGIS) (https://www.qgis.org/en/site/), and the orthophoto obtained from the first flight was used as a reference for geographic calibration. Each plot was cropped based on the shapefile in QGIS. The NDVI map in the first flight was segmented with a threshold of 0.6 for removing background. Finally, four SIs including NDRE, NDVI, GNDVI, and OSAVI were extracted and their specific definitions are provided in Table S2 (Cao et al., 2021) The SIs were indicators of the total quantity and quality of crop photosynthetic activity and were sensitive to variation in canopy chlorophyll concentration, green biomass, and green leaf area(Thompson et al., 2019, Hassan et al., 2019, Gitelson & Merzlyak, 1998, Rondeaux et al., 1996).

### 2.5 Quantification of SG traits

The relative stay green score (RSGS) of individual genotypes to quantify multistage SG trait, at four senescence stages were calculated using temporal indices. These four stages were the milk-ripe stage (S1) (Feekes scale (FS) 11.1), transitional stage from milk-ripe to mealy ripe stage (S2) (FS 11.1 to 11.2), middle mealy ripe stage (S3) (FS 11.2), and late mealy ripe stage (S4) (FS 11.3). The stay green status at anthesis was used as the maximum SG reference (scored as 100), and the relative senescence score (RSS) during a S*i* (*i* = 1, 2, 3, 4) stage (denoted by RSS_S*i*_) was calculated as in formula 2-1. The RSGS at S*i* (*i* = 1, 2, 3, 4) stage (denoted by RSS_S*i*_) was calculated as in formula 2-2 after flowering of all samples.

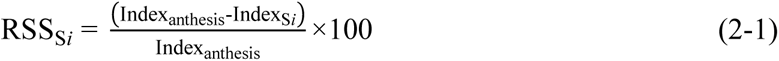

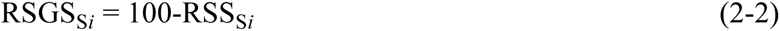

where, Index_anthesis_ and Index_S*i*_ were sample index values at anthesis and a specific S*i* stage, respectively.

The florescence values showed differences among genotypes as early flowering (EF), middle flowering (MF), and late flowering (LF) samples were included in the study. The post-anthesis accumulated temperatures (AT) (sum of daily mean temperatures) of samples were used to enhance the phenotype comparability of EF, LF, and MF members as post-anthesis senescence in wheat is almost exclusively sensitive to temperature *per se* (Reynolds, 2012). Enhancement of phenotypic comparability was performed according to formula 2-3 and multistage AT of EF, LF, and MF samples during the two growing seasons are provided in Table S3.

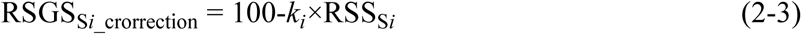

where, *k_i_* is the AT correction factor for sample florescence inconsistency at stage S*_i_*; *k*i is equal to 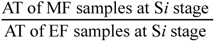 for EF samples and the *k_i_* is equal to 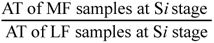 for the LF samples. In addition, the RSGS of MF samples and corrected RSGS of EF and LF samples calculated by each index at S*i* stage were labeled as SG_index*i*_ to facilitate subsequent descriptions.

### 2.6 Heritability analysis

Broad-sense heritability (*H*^2^) was calculated as formula 2-4.

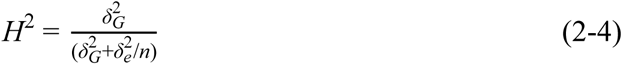

Where, 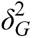 is the genetic variance, 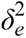 is the residual error variance, n is the number of environments. The values of 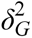 and 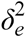 were estimated by analysis of variance (ANOVA) using the lmer function in the lme4 package in the R environment http://www.R-project.org/. The Pearson correlation coefficients (PCCs) analysis was performed using the OmicShare tools (https://www.omicshare.com/tools).

### 2.7 Population structure, PCA, and LD analysis

Population structure was assessed using unlinked markers (*r*^2^ = 0) in STURCTURE 2.3.4 (Earl & vonHoldt, 2011). The model was applied without using prior population information, and the most likely number of subpopulations was determined using a previously described method (Earl & vonHoldt, 2011). Principle component analysis (PCA) was conducted with the GCTA software used for assessing population structure (Yang et al., 2011). Linkage disequilibrium (LD) analysis for the whole genome and A, B, and D genomes used PLINK software (Wu et al., 2021). LD decay distance was obtained by constructing *r*^2^ values against SNP loci and fitting these data to a locally weighted polynomial regression (LOESS) curve using the R program (http://www.R-project.org/).

### 2.8 GWAS, Fst and XP-CLR analysis

GWAS was performed with GEMMA software using a univariate linear mixed model (MLM) (Zhou & Stephens, 2012). The suggested threshold for *P* values ranged from 1.38×10^-4^ to 6.02×10^-4^ for each chromosome and the uniform value *P*=2.60×10^-4^ was adopted as the threshold criterion for genome-wide significance. Continuously significant markers within a distance of 5 Mb were considered a QTL. Identified GWAS loci were compared with previously identified QTL based on their physical locations in Chinese Spring reference genome v2.1. For previously reported SG genes/QTL, the closest flanking markers were used to generate confidence intervals. Whether the loci identified in the GWAS were novel depended on the haplotype block interval.

The population differentiation fixation index (*F*st) and cross-population composite likelihood ratio (XP-CLR) score between wheat genotypes released pre-and post-1970 were calculated for each 100-kb window across the entire wheat genome by VCFtools software (v.4.0), and parameters in the program were set as follows: “--fst -window-size 1,0000,00 --fst-window-step 100,000 --weir-fst-pop” and “--size 1,000 --step 2,000 --ld 0.95”. To detect genes was under selection, the ranking of selection sweeps was based on decreasing *F*st and XP-CLR scores, and the top 5% of regions were chosen as selective sweeps.

### 2.9 RNA-seq

The RNA-seq experiment included mock and heat stress flag leaves of Shaanong 253 and Shanong 15. When wheats grow to 10 days after anthesis, the mock group was treated with normal growth condition of 22℃, 16 hours and 18℃, 8 hours in the greenhouse; while the test group was treated with heat stress condition of 38℃, 8 hours and 24℃, 16 hours in the following three days. Then flag leaves tissues were collected from Shaanong 253 and Shanong 15 at 0, 24, 48, and 72 hours using three independent biological replicates.

Leaf tissues from three biological replicates were combined in equal amounts and send to BGI Genomics, BGI-SHENZHEN (https://www.genomics.cn/) for RNA isolation, RNA library construction and RNA-seq. Quality control, alignment, counting fragments, TPM (transcripts per million Kb calculation) analysis for RNA-seq data as referenced in Yi et al. (Ke Yi, 2023).

### 2.10 Identification of candidate genes

Based on IWGSC RefSeq v2.1 gene annotations (http://wheatomics.sdau.edu.cn/), high confidence (HC) genes located within QTL were used for candidate gene analysis. Candidate genes or homologs amino acid sequences for SG in other plant species were downloaded from https://bigd.big.ac.cn/lsd/ to compare with potential wheat genes (E value <10-5). Gene Ontology (GO) functional enrichment was performed using http://wheat.cau.edu.cn/TGT/ (Chen et al., 2020) and visualization by clusterProfiler (v3.18.1) in the R package. Genes in the senescence pathway were selected as potential candidate genes. Simultaneously, the candidate genes were further confirmed by the expression levels. Finally, candidate gene-based association analysis was conducted using the Mixed Linear Model (MLM) method in GAPIT 3.0 and *P* = 1×10-^4^ as the threshold for significance.

## 3. Results

### 3.1 Genetic diversity of wheat panel

Population structure and principal components analysis (PCA) based on Bayesian clustering identified six sub-populations (Sp) (Fig. 2B). LD (r^2^) analyzed by pairwise comparisons of 361,293 SNP decayed to the critical r^2^ value (0.1) was estimated at about 3.8 Mb for the whole genome (Fig. 2C); LD decay of 1.8 Mb was faster for the D genome followed by the A and B genomes at 3.1 Mb and 6.1 Mb, respectively (Fig. 2C). The faster LD decay in the D genome was related to relatively less artificial selection after incorporation into common wheat. In population structure, Sp1 (Fig. 2A) consisted mainly of Chinese landraces (CL); Sp2 mainly contained introduced modern cultivars (IMC); Sp3 mainly included Chinese spring cultivars (CSC); Sp4 predominantly included mixed winter cultivars (MWC); Sp5 comprised Chinese winter cultivars, mostly pre-2000 (CHWC, pre-2000; and Sp6 mainly included Chinese winter cultivars grown after 2000 (CHWC, post-2000). Sp3-Sp6 were combined as modern Chinese cultivars (MCC) (Fig. 2A). In the process of senescence, CL, IMC, and MMC exhibited significant differences in SG phenotype (Fig. 2D and 2E). Geographical origin, historical timing and temperature/light characteristics were major factors that determined the diversity groups in this panel.

**Fig. 2.**
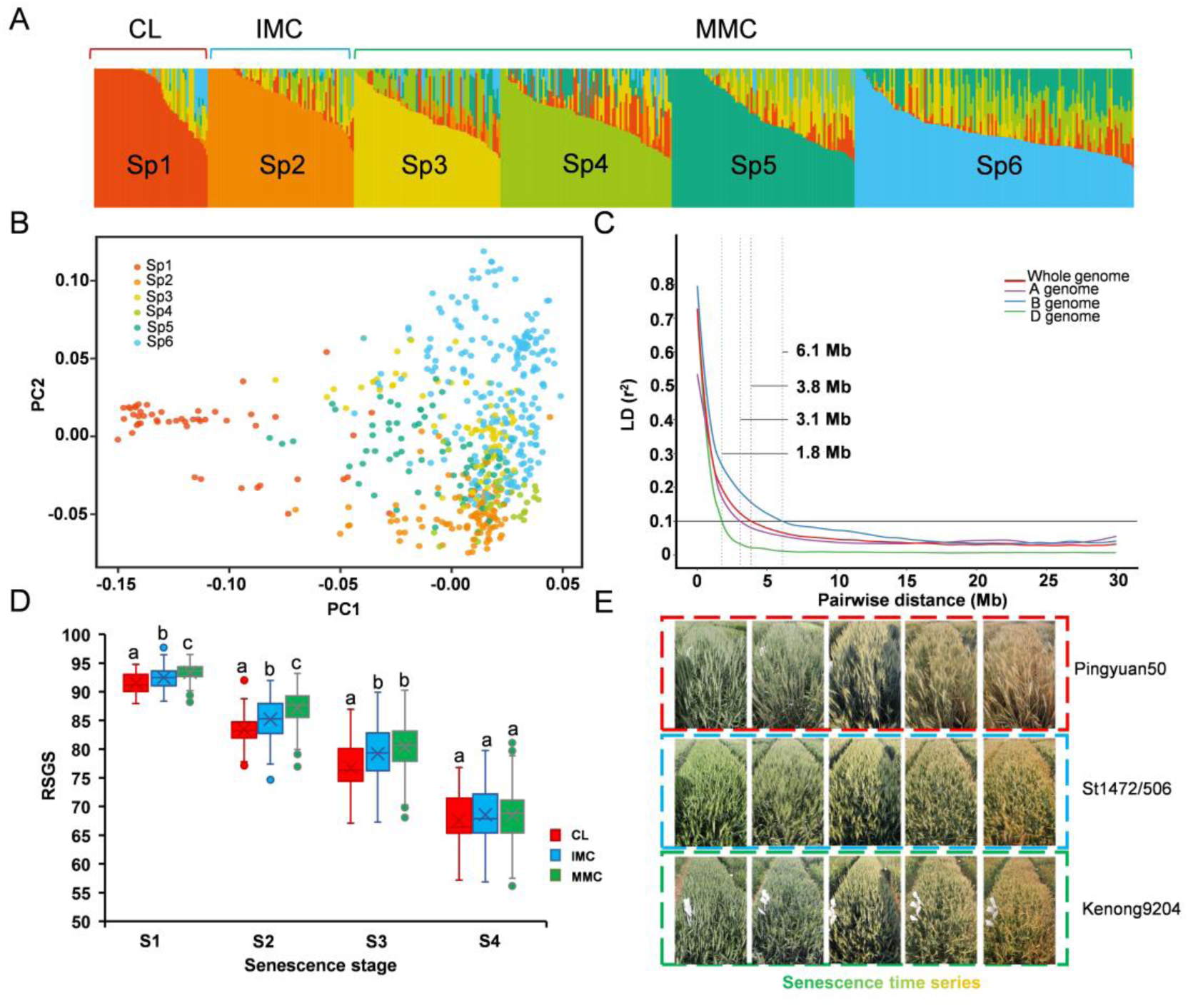
Genetic structure of wheat panels. (A) Population structure of wheat panels. (B) Principal coordinates analysis (PCA) of wheat panels. (C) Linkage disequilibrium (LD) decay over physical distances. Plots of pair-wise single-nucleotide polymorphism LD (r^2^) values as a function of inter-marker map distance (Mb) within three subgenomes. (D) SG score of CL, IMC, and MMC based on SGGNDVI in 2022 indicate the significant differences (*p* < 0.05, LSD test). (E) The SG phenotype of Pingyuan 50 (CL), St1472/506 (IMC), and Kenong 9204 (MCC) in the senescence time series.

### 3.2 Evaluation of phenotypic performance

The average dynamics of the SIs of all genotypes during the 2020-2021 and 2021-2022 seasons are shown in Fig. 3. All SI values showed a consistent decline over post-anthesis thermal time. The decline was initially but increased with approaching physiological maturity. The decline was relatively slow during the watery ripe and milky ripe stage stages (post-anthesis AT <400 ℃ per day) as there was less competition for assimilates from source to sink, but was relatively fast from the mealy ripe stage (post-anthesis AT >400 ℃ꞏper day) as competition for assimilates increased. These temporal SIs robustly tracked the senescence process in the wheat canopy in field environments.

**Fig. 3.**
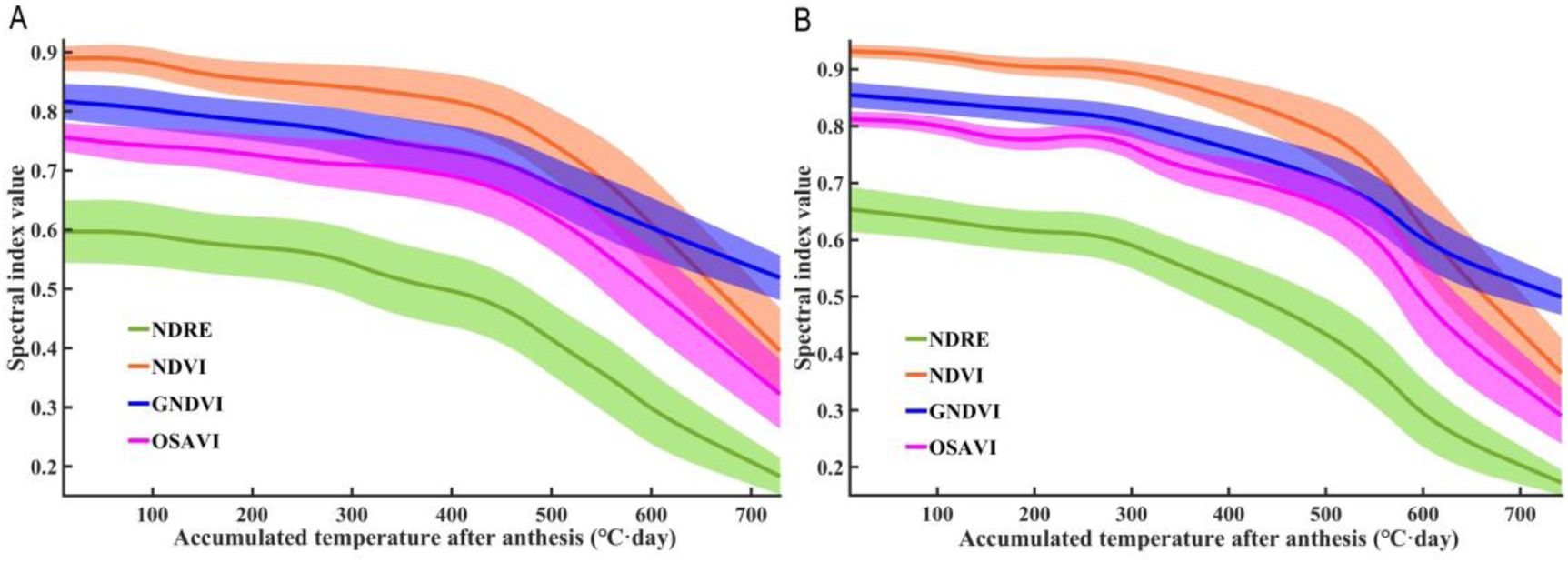
Dynamics of fitted temporal SIs in 2020-2021 (A) and 2021-2022 (B). Lines represent index mean values and shaded areas represent standard deviations (sd).

The quantitative results of SG traits for the wheat genotypes during both seasons are shown in Table 2. The mean values for the SG_index_ decreased over post-anthesis thermal time and the standard deviations increased. The numerical range of each index varied from stage S1 to S4. For S1 and S2, the values of SG_NDRE_ were lower than the those of SG_NDVI_ and SG_OSAVI_ in both years and temporal NDRE revealed greater relative senescence scores (>12 at S1 and >18 at S2) for the canopy during the early senescence phase in both years. The PCCs between different RSGS traits during both seasons are shown in Fig. S2. For RSGS calculated using each index, there was a generally strong linear correlation (PCCs >0.82) between SG_index1_ and SG_index2_, but the linear relationships of SG_index3_ with SG_index2_ and SG_index4_ were also strong (PCCs >0.83) in both years. The PCCs of RSGS were high (0.17∼0.73) between the two cropping seasons.

**Table 2.**
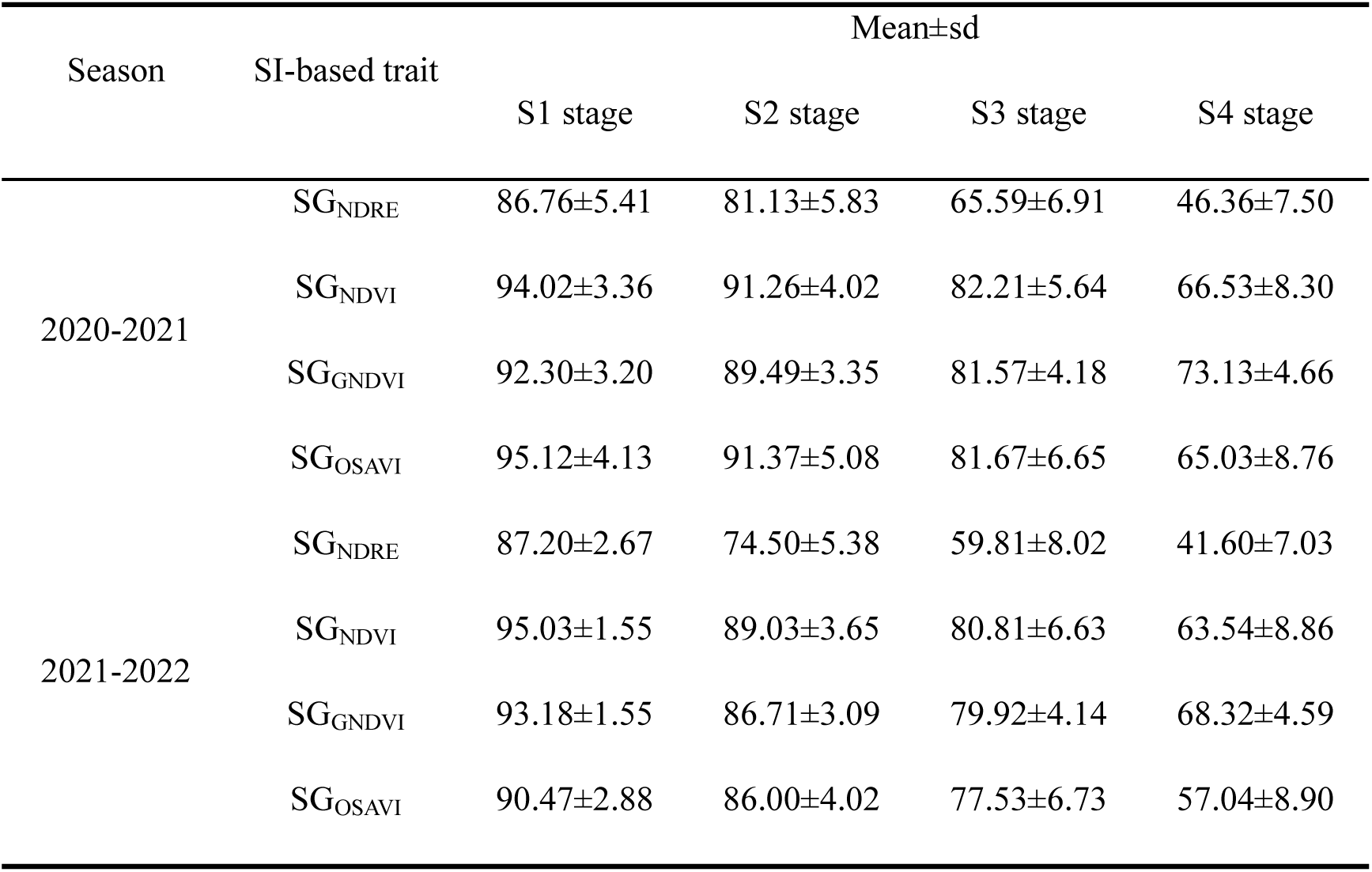
Means and standard deviations of SI-based traits in different seasons.

The *H*^2^ of all SG traits among the 4 stages ranged from 0.31 to 0.84, with an average of 0.65 (Table S4). Among the 4 SG traits, heritability increases with the increase of senescence stage, while SG_OSAVI_ exhibits higher heritability than other SG traits in the 4 stages. Moreover, RSGS in CL, IMC, and MMC, including varieties released pre- and post-1970 showed significant differences (Fig 1D and S3). MMC had a higher RSGS compared to IMC and CL, and varieties released post-1970 had a higher RSGS than those released in pre-1970.

### 3.3 GWAS of SI-based traits

GWAS performed to explore the genetic basis of SG-related QTL in the diversity panel detected 47 high-confidence QTL harboring 3,079 significant SNPs associated with the SI-based traits during the two years (Table S5). The R^2^ by each QTL ranged from 0.1% to 26%, with a mean of 3.4% (Table S5). The chromosomal distribution of QTL is shown in Fig. 4 and Manhattan plots of the SI-based traits are listed in Fig. S4 to S7. Thirty and 32 QTL were detected in 2020-2021 and 2021-2022, respectively (Table S5; Fig. S8) with 15 being co-located in both seasons (Table S5). Fifteen QTL overlapped or were adjacent to previously reported QTL or genes (Table 3; Fig. 4). For example, *Qsg.nwafu-5BL.3* detected on Chr 5B by four SG_index_ in 2020-2021 and 2021-2022 overlapped the reported SG-related QTL *QSg.sau-5B.3* (Ren et al., 2022). The SG-related gene *TaARF15* in *Qsg.nwafu-6BL.2* was detected using SG_OSAVI_ in 2020-2021 and SG_NDVI_, SG_OSAVI_, and SG_NDRE_ in 2021-2022(Li et al., 2023). The number of QTL detected by each SI-based trait during the entire senescence stage differed between years. In 2020-2021 the most QTL were identified by SG_GNDVI_, followed by SG_NDVI_, SG_OSAVI_, and SG_NDRE_ (Fig. S8); 2021-2022 most the most QTL were identified by SG_NDRE_, followed by SG_OSAVI_, SG_NDVI_, and SG_GNDVI_ (Fig. S8). Moreover, the numbers of QTL detected by the SI-based traits differed at multiple senescence stages. During the two seasons, 7 QTL were located only at stages S1 and S2, and 4 were located only at S3 and S4; 36 QTL were located at least three out of four stages (Table S5).

**Fig. 4.**
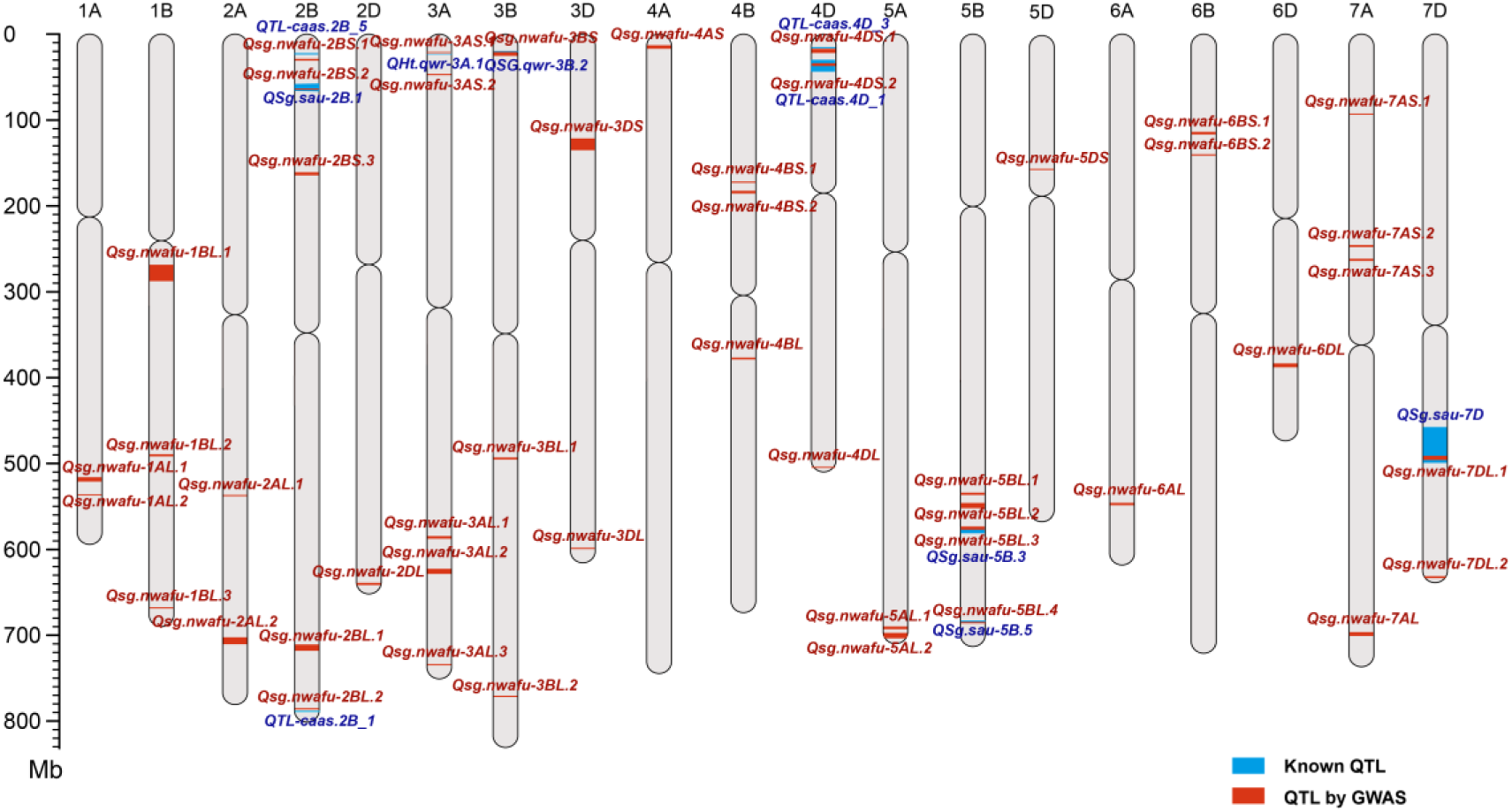
Distribution of the SG QTL from this study and known SG QTL on wheat chromosomes. Blue bands represent known SG QTL; red bands represent SG QTL detected by GWAS in the present study; band widths represent the confidence intervals.

**Table 3.**
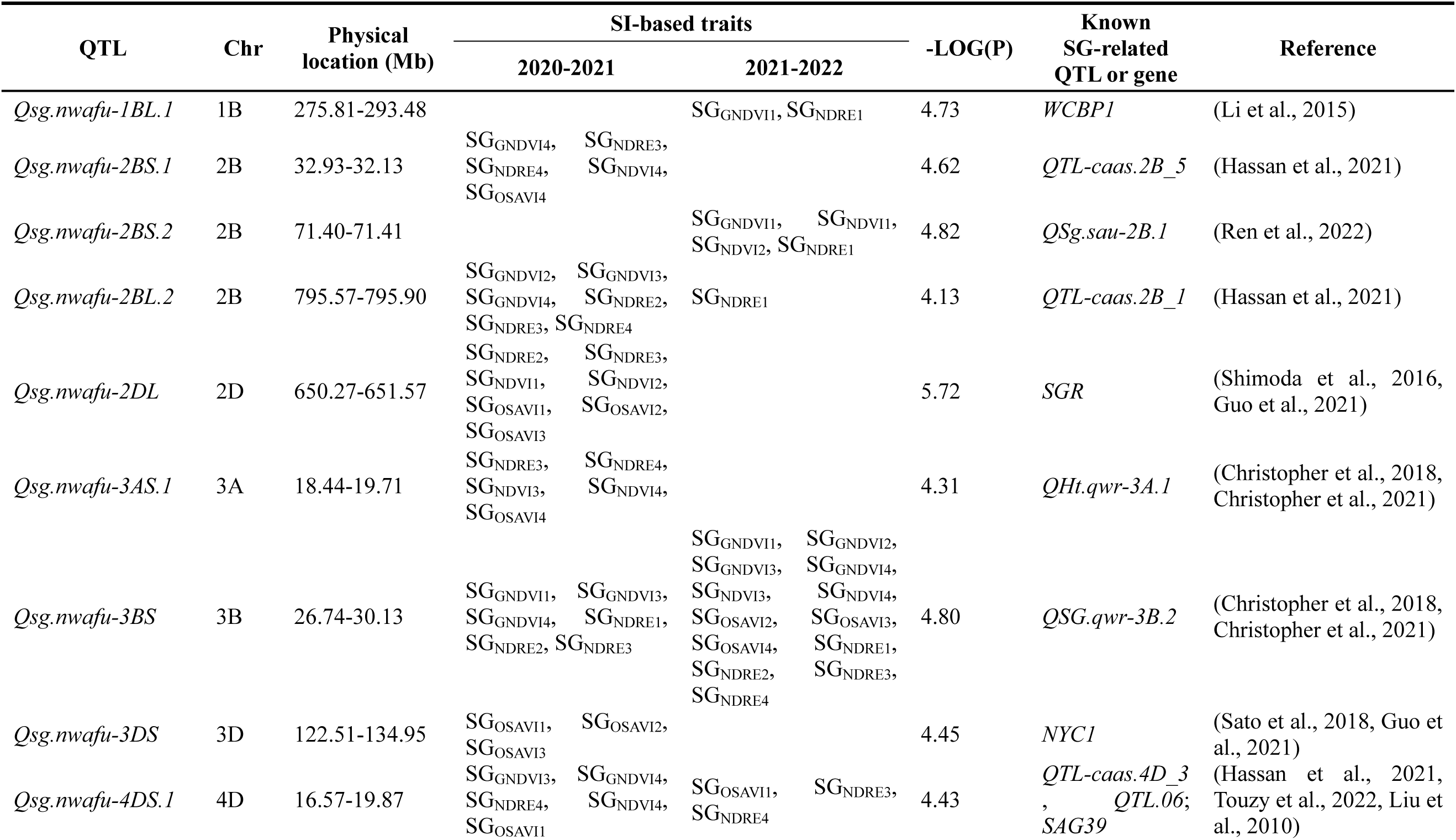
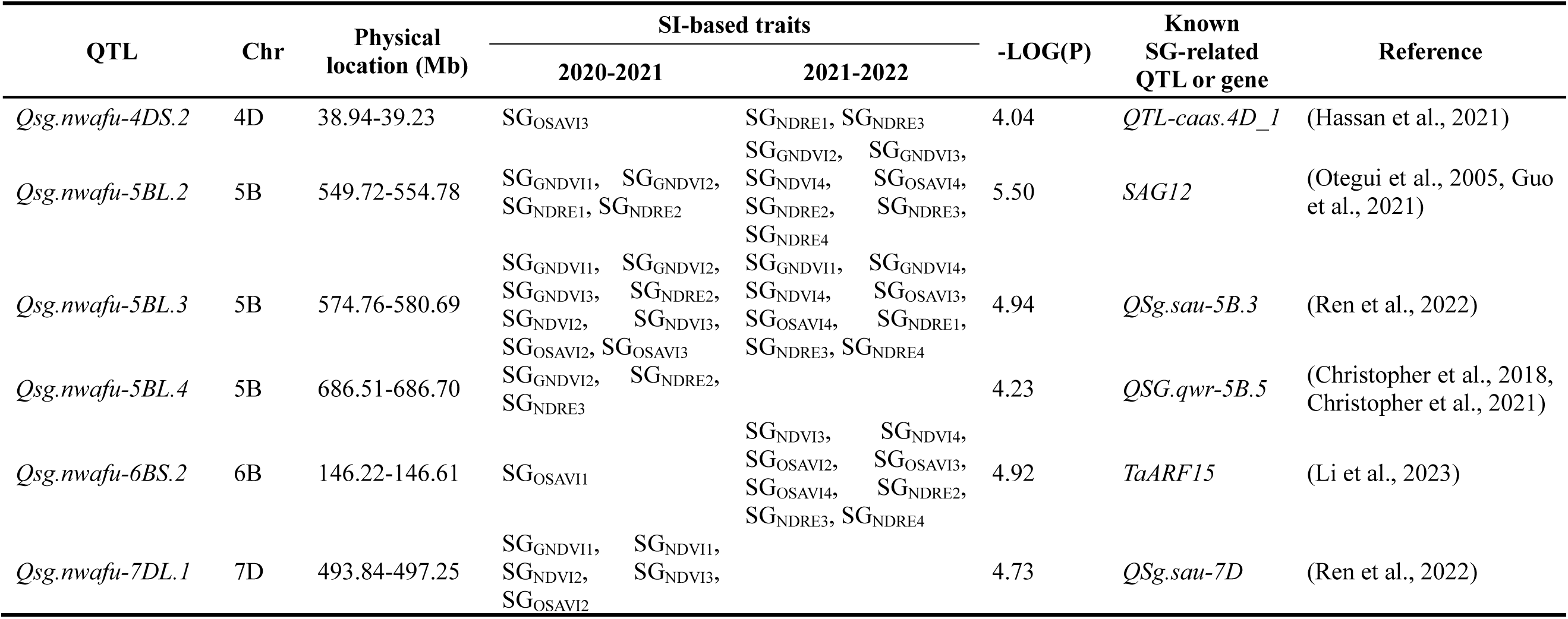
QTL overlapping or adjacent to reported SG-related QTL or genes.

### 3.4 Candidate genes for SG-related traits

We complied with three principles for candidate genes analysis in the confidence interval: 1) if the QTL region contained known genes controlling SG, those genes were considered high-priority candidate genes; 2) if the QTL regions contained wheat homologs associated with genes regulating SG or senescence in other species, those genes were considered priority candidate genes; 3) if the QTL regions did not contain either of these types the QTL were considered to be new loci.

Based on the three principles for candidate genes analysis, we identified a number of potential causal genes. For example, *TaARF15* (*TraesCS6B03G0356400*) was identified in *Qsg.nwafu-6BS.2* and has been to be involved in the JA mediated wheat senescence pathway in recent research(Li et al., 2023). *TraesCS2A03G1081100* (*D2HGDH*) in *Qsg.nwafu-2AL.2* was identified as a candidate gene through homologous alignment of a SG protein sequence in Arabidopsis. There was 72.43% homology with *AT4G36400* (*D2HGDH*) at the protein level. *AT4G36400* is in the same network as several genes involved in β-oxidation and degradation of branched-chain amino acids in chlorophyll that delays senescence to some extent (Engqvist et al., 2009). In addition, GO enrichment analysis of 1,085 high-confidence candidate genes located in the QTL showed that some of the genes were directly or indirectly involvement in plant senescence. We found 37 GO terms (p <0.05) among which the top 35 of the three GO enrichment types are shown in Table S6 and Fig S9. This analysis strongly suggested that these genes are significantly associated with senescence-associated vacuoles in cellular components; and leaf senescence, ethylene response, and programmed cell death in biological processes. The functions of these genes are mostly associated with nutrient reservoir activity. In biological processes, *TraesCS2B03G1299500* (*WRKY70*) in *Qsg.nwafu-2BL.1* belongs to GO:0010150 and participates in leaf senescence downstream of the developmental process pathways. Based on *WRKY70* as a negative regulator of leaf senescence in Arabidopsis (Besseau et al., 2012), we considered it as a key candidate gene for SG. Moreover, all of the three candidate genes showed increased expression levels in Shaannong 253 (SG) compared with those in Shannong 15 (Non SG) at different time points after heat stress (Fig. S10).

### 3.5 Selection of favorable SG haplotypes in breeding

The reported gene *TaARF15-B1* (*TraesCS6B03G0356400*) and priority genes *TraesCS2B03G1299500* (*WRKY70*) and *TraesCS2A03G1081100* (*D2HGDH*) identified in this study were selected for validation in breeding. To detect indirect selection of SG-related loci or genes during modern breeding we divided the panels into pre- and post-1970 groups (Fig. S3). Then, *F*st and XP-CLR was used to detect putatively selected regions during modern breeding.

*F*st and XP-CLR analyses of *TaARF15-B1* located in *Qsg.nwafu-6BS.2* showed significant signals in the QTL region (Fig. 5A). In GWAS two significant SNPs, *s6B146446687* and *s6B146446689*, located at 2388 bp (T/A, Glu/Asp) and 2390 bp (C/A, Glu/Stop codon) in the *TaARF15-B1* coding sequence caused a missense and a termination of transcription, respectively (Fig. 5B). The effects of the Hap1 (A-A) and Hap2 (T-C) variations on the dynamic SI-based traits, thousand kernel weight (TKW), yield and crude protein were determined. Hap1 (186 accessions) had significantly higher SI-based traits, TKW and yield than Hap2 (56 accessions), but the crude protein showed an opposite significant difference (Fig. 5C to F). Furthermore, the Hap1 and Hap2 were explored the history of modern breeding in 805 worldwide wheat accessions (Table S7). The frequency of Hap1 increased from 29% pre-1951 to 57.5% 1951-1970 among Chinese wheat accessions and reached 90% in the post-2010 (Fig. 5G). Concurrently, we determined that the frequencies the favorable Hap 1 in wheat zones Ⅰ, Ⅱ, Ⅲ, Ⅳ, Ⅵ, and VIII were relatively high at 61%, 81%, 60%, 57%, 54% and 52%, respectively (Fig. 5H).

**Fig. 5.**
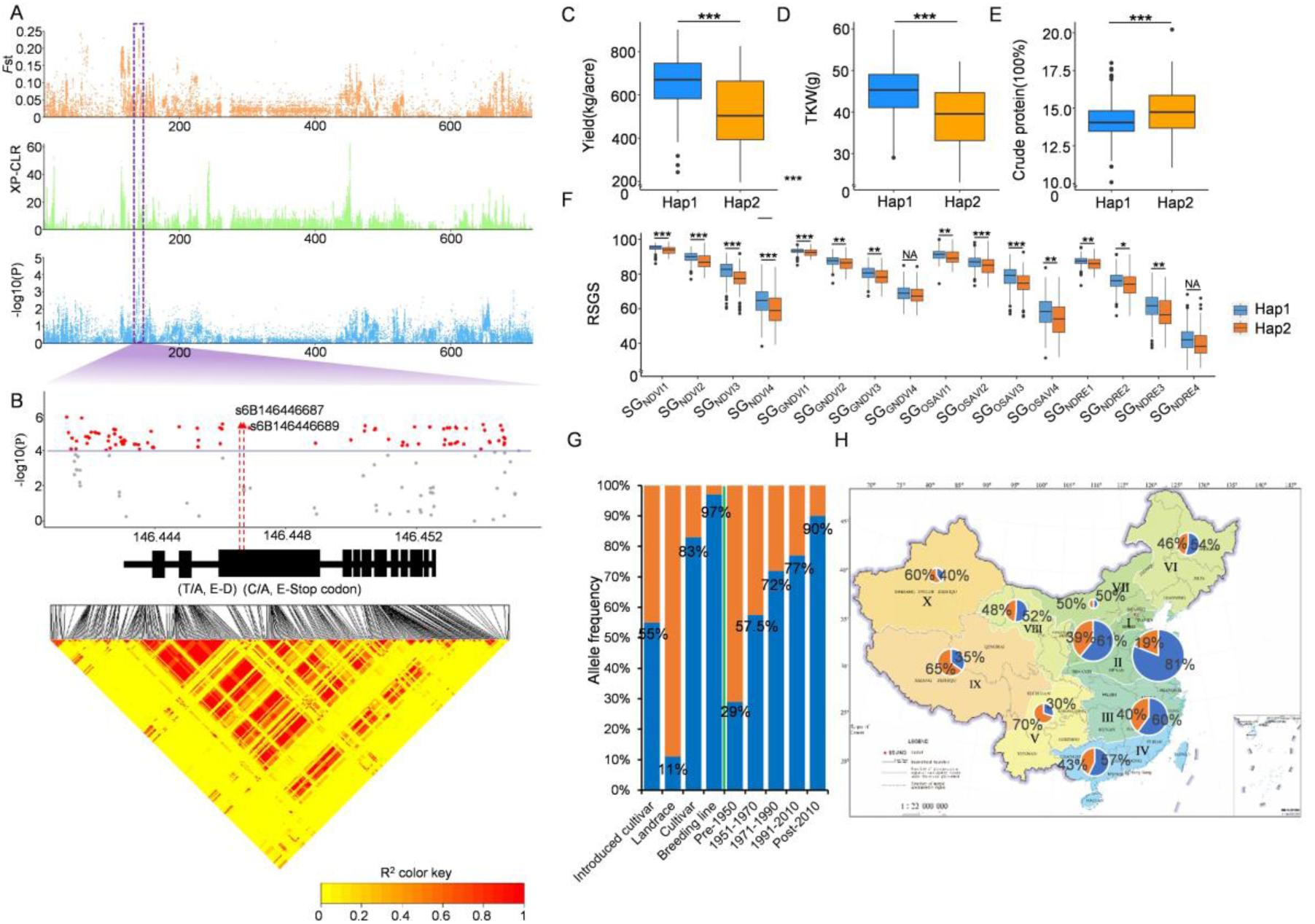
Variation in *TraesCS6B03G0356400*. (A) *F*st, XP-CLR, and associated significant signals on chromosome 6B. Purple rectangle is the target associated signal position on 6B and purple shading is the significance threshold (−log10[*P*-value] =4.0). (B) Local Manhattan plot (top) and LD heat map (bottom) surrounding *TraesCS6B03G0356400*. Red color indicates strong LD with the significant variation. Two red triangles represent two variations in *TraesCS6B03G0356400*. (C-F) Mean yield, thousand kernel weight, crude protein and SG scores for the two haplotypes. *p* values were calculated using two-tailed t-tests (*, *P* <0.05; **, *P* <0.01; ***, *P* <0.001; NS, not significant). (G) Percentages of haplotypes in different wheat categories (left of the green line) and breeding periods (right of the green line (Hap1, blue; Hap2, orange). (H) Frequencies of haplotypes in different wheat zones. Circle size represents population size; Hap1, blue; Hap2, orange).

There was also a relatively high selection signal in *Qsg.nwafu-2AL.2* (Fig. 6A). Three salient SNPs *s2A709642415*, *s2A709643989* and *s2A709645459* were identified in *TraesCS2A03G1081100* (*D2HGDH*); *s2A709645459* caused a synonymous variation at 410 bp in the gene, and *s2A709642415* and *s2A709643989* caused missense variations at 135 bp (G/C, Gly/Arg) and 252 (T/C, Phe/Ser) bp in the coding sequence (Fig. 6B). Dynamic stay green, thousand-kernel weight, and yield showed significant differences between the Hap1 (C-C) group (86 accessions) and Hap2 (G-T) group (158 accessions) (Fig. 6C, D and F) but there was no significant difference in crude protein content (Fig. 6E). Hap1 was identified as the favorable haplotype. The history of the two major haplotypes, Hap1 (195 accessions) and Hap2 (603 accessions) in the 798 worldwide wheat accessions released before 2020 indicated that Hap1 increased in frequency from 7.5% before 1950 to 47% post-201 this indicating strong selection (Fig. 6G; Table S8). The frequency of Hap1 in zones Ⅰ, Ⅱ and Ⅷ were 39%, 40% and 19%, respectively (Fig. 6H; Table S8).

**Fig. 6.**
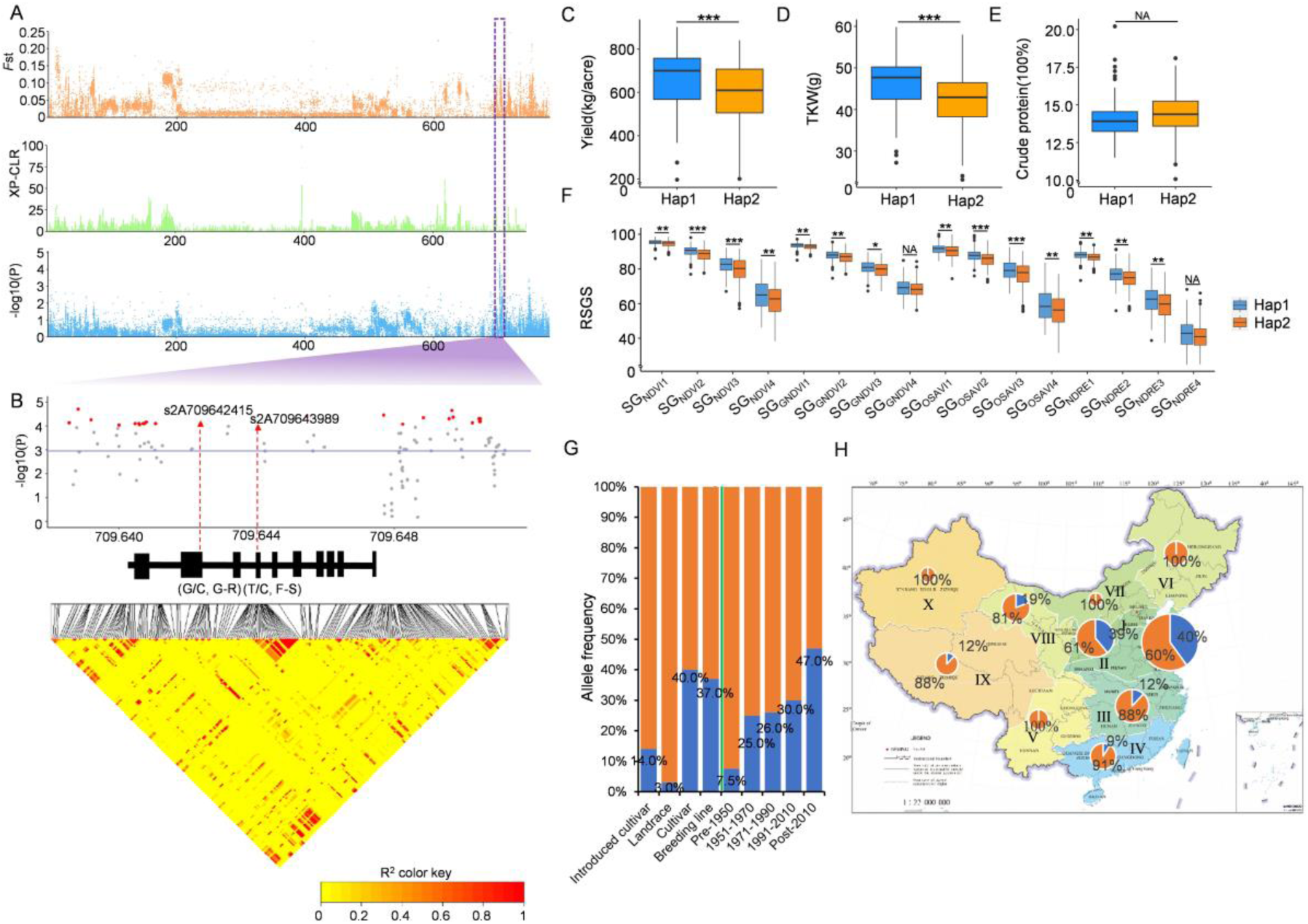
Variation in *TraesCS2A03G1081100*. (A) The *F*st, XP-CLR, and associated significant signals on 2A. Purple rectangle is the associated signal position in chromosome 2A and purple shading shows the significance threshold (−log10[*P*-value] =4.0). (B) Local Manhattan plot (top) and LD heat map (bottom) surrounding *TraesCS2A03G1081100*. Red color highlights strong LD with significant variation. Red triangles represent two variations in *TraesCS6B03G0356400*. (C-F) Thousand kernel weight, crude protein and canopy greenness scores for the two haplotypes. *p* values were calculated using two-tailed t-tests (*, *P* <0.05; **, *P* <0.01; ***, *P* <0.001; NS, not significant). (G) The percentages of the two haplotypes among wheat groups (left of the green line) and breeding periods (right of the green line) (Hap1, blue; Hap2, yellow). (H) Percentages of the two haplotypes in different wheat zones. Circle size represents the number of accessions; Hap1, blue; Hap2, orange).

*F*st and XP-CLR of the *Qsg.nwafu-2BL.1* region showed significant signals (Fig. 7A). The salient SNP *s2B720780102* caused a missense variation at 177 bp (G/A, Gly/Ser) bp in the *TraesCS2B03G1299500* (*WRKY70*) coding sequence (Fig. 7B). Hap1 (G) and Hap2 (A) were compared to determine the effects of the dynamic SI-based traits, thousand kernel weight (TKW), yield and crude protein. The Hap1 group (199 accessions) had significantly higher SI-based traits, TKW and yield than the Hap2 group (47 accessions), and crude protein again showed an opposite significant difference (Fig. 7C to F). The history of Hap1 (787 accessions) and Hap2 (206 accessions) in Chinese wheat accessions released before 2020 indicated that Hap1 increased in frequency from 46% of accessions pre-1950 to 93% post-2010 (Fig. 7G; Table S9). The frequency of Hap 1 in wheat zones Ⅰ, Ⅱ, Ⅲ, Ⅳ, and Ⅴ were 85%, 66%, 74%, and 62.5%, respectively (Fig. 7H; Table S9).

**Fig. 7.**
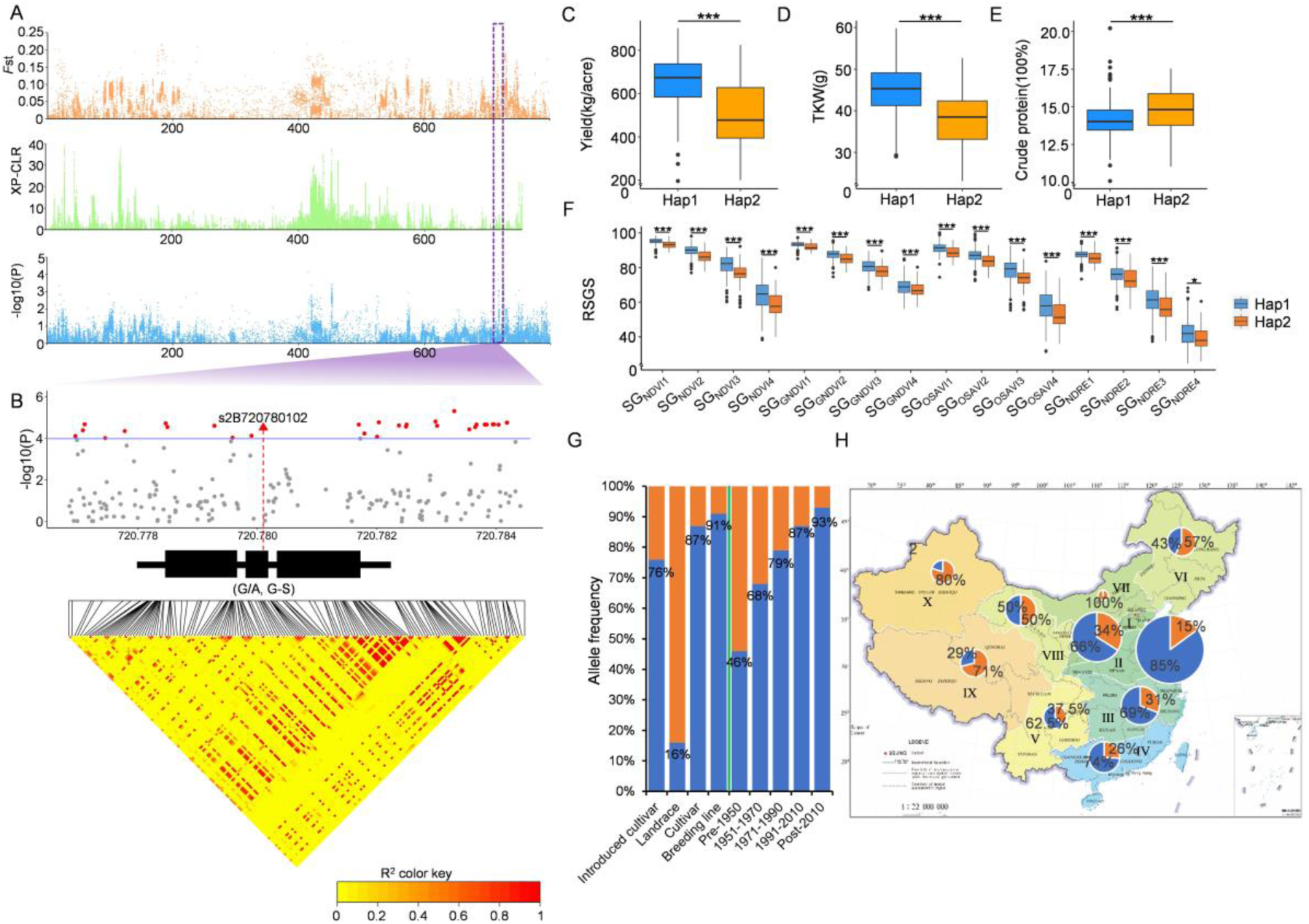
Variation in *TraesCS2B03G1299500*. (A) The *F*st, XP-CLR, and associated significant signals on chromosome 2B. Purple rectange indicates the target associated signal position on chrosome 2B; purple shading is the significance threshold (−log10[*P*-value] = 4.0). (B) Local Manhattan plot (top) and LD heat map (bottom) surrounding *TraesCS2B03G1299500*. Red color highlights strong LD with the SNP. Red triangle represents one variant in *TraesCS6B03G0356400*. (C-F) The distribution of the two haplotypes corresponding length of Mean yield, thousand kernel weight, crude protein and canopy greenness scores for two haplotypes. *p* values were calculated using two-tailed t-tests (*, *P* <0.05; **, *P* <0.01; ***, *P* <0.001; NS, not significant). (G) Frerquencies (%) of two haplotypes in different categories (left of the green line) and breeding periods (right of the green line (Hap1, blue; Hap2, yellow). (H) The percentages of two haplotypes in different wheat zones. Circle size indicates the number of cultivars; (Hap1, blue; Hap2, yellow).

Along with the higher frequencies of favorable SG haplotypes in modern Chinese cultivars and breeding lines the frequencies in introduced cultivars were much higher than in the CL group (Fig. 5G, Fig. 6G, and Fig. 7G).

## 4. Discussion

### 4.1 High-throughput phenotyping of SG traits in the wheat diversity panel

For this study we attempted to select diverse accession panels with relatively concentrated flowering periods and excluded lines on the basis of lodging and disease susceptibility. Analysis of genetic diversity, population structure, and PCA confirmed that the panels had sufficient diversity, and the trends in LD decay were consistent with previous research (Yu et al., 2020, Wu et al., 2021).

Stay green in wheat is a complex dynamic trait. Senescence in wheat starts from the lower leaves and progresses to the flag leaves, whereas the senescence of a single leaf generally starts from the tip and progresses to the base (Kamal et al., 2019, Sultana et al., 2021). SG assessment was more effective at the canopy level in the wheat diversity panel in the field. Reflectance SIs derived from imaging or non-imaging sensors provide unique opportunities for acquiring data on the dynamics of wheat canopy senescence from large populations (Inoue et al., 2008). NDVI obtained from handheld, multispectral, or hyperspectral sensors are widely used for crop senescence detection and show potential in determining the senescence patterns in wheat genotypes (Christopher et al., 2021, Pinto et al., 2016, Christopher et al., 2016). In the present work, the dynamics of NDVI during wheat senescence in diversity panel were consistent with previous observations. Hassan, M., et al. showed that NDRE and GNDVI were also reliable in revealing the spatiotemporal senescence dynamics of multiple wheat accessions in the field(Hassan et al., 2018). NDRE showed stronger correlations with wheat chlorophyll content and leaf area index (Thompson et al., 2019). GNDVI, measures the green spectrum instead of the red spectrum, and is similar to NDVI and more sensitive to chlorophyll concentration (Gitelson & Merzlyak, 1998). In this study, NDRE showed a lower saturation effect during early senescence. NDRE is calculated by including the red edge band and the red edge spectrum (690-740 nm) in the crop reflectance curve thereby including transition from chlorophyll absorption and near-infrared leaf scattering (Thompson et al., 2019). In addition, the high heritability of RSGS calculated from the 4 SIs reflects effective explanatory power for the SG phenotype. Our results showed that NDRE, NDVI, GNDVI, and OSAVI could reveal canopy senescence of wheat populations in complex field situations.

Effective SG quantification in wheat populations is essential to identify novel genes. Previous studies proposed useful SG quantification pipelines for SG-related QTL mapping in wheat populations using temporal SIs(Hassan et al., 2021, Christopher et al., 2021). In this study, a more targeted pipeline of SG quantification using temporal SIs was proposed for diverse wheat panels. This pipeline calculated RSGS for SG quantification and RSGS can be interpreted as the proportion of canopy greenness of a genotype during the post-anthesis stages. In other words, RGRS characterizes the relative degree of delayed senescence. Moreover, the post-anthesis AT of samples were incorporated to enhance the comparability of phenotypic data to deal with inconsistency of samples florescence. The SG traits quantified by the pipeline were efficient in revealing multi-stage SG differences between high stay green and low stay green varieties. The pipeline was therefore considered useful for GWAS of wheat diversity panels.

### 4.2 SG QTL detection in the wheat diversity panel

In order to mine SG genes it was necessary to trace and accurately quantify dynamic changes in stay green related traits. Previous studies proposed useful quantification frameworks for mapping QTL underlying SG in RIL, DH and MR-NAM populations using temporal SIs. For example, Christopher et al. (Christopher et al., 2021, Christopher et al., 2018) fitted logistic functions using temporal NDVI to calculate SG parameters in doubled haploid (DH) and multi-reference nested association mapping (MR-NAM) wheat populations. In those studies, 43 SG-related QTL were identified in DH populations and 65 were identified in MR-NAM populations. Pinto et al. (Pinto et al., 2016) used temporal NDVI to calculate sample SR (slope of the NDVI decline against thermal time), greenness decay (percentage of NDVI decline), and the area under the NDVI curve in recombinant populations of inbred lines (RIL); 44 QTL were detected. Muhammad et al. (Hassan et al., 2021) evaluated multistage SG traits in DH wheat populations using temporal data for six SIs. SG traits were determined by absolute decline of the SIs and were measured by subtracting the SI values at the 10%, 50% and 90% senescence stages from the SI value at heading; 28 senescence-related QTL were identified. Visual senescence score was used to evaluate SG in wheat RIL populations and 40 QTL were mapped (Ren et al., 2022). However, SI-based traits such as single-stage indexes or absolute senescence used in these various studies had a low interpretable ability when comparing differences between individuals in diverse germplasm panels and were therefore not suitable for GWAS of populations with different phenologies. In the present study, we used the RSGS of samples for SG quantification and GWAS. Over the two seasons stable QTL in the diverse wheat panels were detected at multiple senescence stages. Moreover, some of the SG-related QTL overlapped or were close to previously reported genes, such as *TaARF15-B1* (Li et al., 2023), *NYC1*, and *SAG12* (Guo et al., 2021), providing confidence in our results. Others QTL were probably novel. GO enrichment analysis of candidate genes located in the QTL further supported the validity of phenotype and provided guidance for discovery of more SG-related genes in future studies. The SG phenotyping pipeline using GWAS showed greater potential in localizing SG-related QTL compared to earlier methods.

The work also showed that multiple SIs can be combined to compensate potential defects in using a single SI or inter-annual environmental changes. Transcriptomic studies have revealed that senescence in wheat is a genetically programmed process and occurs in a temporally coordinated mode (Borrill et al., 2019). In the present study, some QTL were detected only during the slow senescence phase or the fast senescence phase. Therefore, phenotyping data from multiple stages of senescence appears to be essential for complete analysis.

### 4.3 Utilization of SG variations in modern breeding

The genetic improvement of wheat has been subject to the influence of human domestication and selection. Discovery and use of favorable SG-related variation in wheat is necessary for developing cultivars with superior SG attributes. We found that wheat germplasm variable in SG traits prior to, and after, 1970, particularly at the S4 phase. Selection signals indicated that the frequency of some key SG-related genes or QTL had gradually increased after 1970, especially in production zones Ⅰ, Ⅱ, and Ⅲ. For example, *TraesCS6B03G0356400* Hap1 associated with higher yield and stay green underwent a clear increase during 1951-1970, the period of the Green Revolution when new varieties were developed from introduced germplasm or crosses of local and foreign genotypes(Thomas & Ougham, 2014, Chapman et al., 2021).

SG-related favorable variations also exhibit unique patterns in given regions. In this study, the frequency of favorable SG-related haplotypes in humid and semi-humid area was higher than arid and semi-arid regions, especially in the Ⅱ and Ⅲ wheat regions. For example, the frequency of favorable SG-related haplotype Hap1(G) in WRKY70 reached 85% and 69% in the Ⅱ and Ⅲ wheat regions, respectively. It is evident that particular ecological wheat regions in China rely heavily on haplotypes that carry the SG trait to accommodate the demands of cropping system and densely populated regions. In Li, H., et al. work, favorable SG-related haplotype in wheat has a similar pattern, the selection of early maturing haplotype *TaARF15-A1-HapI* in arid and semi-arid regions (I, VI, VII, and VIII wheat regions) was higher than humid and semi-humid area to match the requirement for two cropping seasons per year. The frequency of SG haplotype *TaARF15-A1-HapⅡ* was higher in Ⅱ, Ⅲ, and Ⅳ wheat regions (Li et al., 2023).

Senescence is a dynamic process that involves well-orchestrated degradation and remobilization processes that affect crop productivity and quality. Previous studies suggest that SG increases grain yield, but negatively affects nitrogen conversion, especially in leguminous crops (Sultana et al., 2021, Miryeganeh, 2021). In wheat, more than 50% of N accumulation in grains is contributed by chlorophyll degradation in flag leaves (Sultana et al., 2021). Stay green reduces the speed of chlorophyll degradation. As senescence progresses, the chloroplasts and chlorophyll in green tissue degrades, and the speed of senescence affects N transport (Woo et al., 2019, Guo et al., 2021). Hence, the inquiry of physiologists and breeders is centered on the manner by which the onset and speed of senescence is regulated to synchronize the correlation between the sink and the source. This synchronization is crucial to ensure that nitrogen accumulation is not adversely impacted while simultaneously enhancing the yield. In this study, *TaARF15-B1* Hap1 and *WRKY70 Hap1* were favorable SG genotypes for high TKW and yield, but the crude protein contents were comparatively low, rendering them more difficult to meet quality requirements for varietal release. However, the favorable *TraesCS2A03G1081100* Hap1 did not affect crude protein content, a feature requiring more investigation. Since this haplotype was present at relatively low frequency in modern germplasm it could be a target for selection.

## 5. Conclusions

The UAV-based spectral imaging platform provides a convenient tool to quickly obtain the canopy spectral indices from the large populations of crop germplasm in the field. This study explored the high-throughput phenotyping pipeline applicable to SG-related QTL discovery in a diverse wheat panel using temporal spectral indices. The temporal data of NDRE, NDVI, GNDVI, and OSAVI could be robustly used for phenotyping the senescence dynamic of diverse wheat populations in the field. The SI-based traits calculated by fusing indices and accumulated temperature data were effective in GWAS for SG-related QTL and candidate genes detection and many rare reported SG-related QTL could be responded. Meanwhile, haplotypes analysis has shown that the SG trait has been strongly selected in wheat breeding in specific wheat regions in China. The framework proposed in this study provides a referable phenotyping solution to accelerate the elucidation of the SG genetic mechanism in wheat from the diverse panels.

## List of acronyms

AT: Accumulated temperatures
CRP: Calibrated reflectance panel
CL: Chinese landraces
CSC: Chinese spring cultivars
CHWC: Chinese winter cultivars
cM: CentiMorgans
XP-CLR: Cross-population composite likelihood ratio
DH: Doubled haploid
EF: Early flowering
FS: Feekes scale
Fst: F-statistics
GO: Gene ontology
GWAS: Genome-wide association studies
GPS: Global position system
GNDVI: Green normalized difference vegetation index
GSD: Ground sample distance
HC: High confidence
IMC: Introduced (foreign) modern cultivars
IWGSC: International Wheat Genome Sequencing Consortium
LF: Late flowering
MLM: Linear mixed model
LD: Linkage disequilibrium
MF: Middle flowering
MWC: Mixed winter cultivars
MCC: Modern Chinese cultivars
MR-NAM: Multi-reference nested association mapping
NDRE: Normalized difference red edge index
NDVI: Normalized difference vegetation index
OSAVI: Optimized soil-adjusted vegetation index
PCCs: Pearson correlation coefficients
PCA: Principle component analysis
QTL: Quantitative trait loci
QGIS: Quantum GIS
RIL: Recombinant inbred lines
RSS: Relative senescence score
RSGS: Relative stay green scores
SIs: Spectral reflectance indices
Si: Stagei
SG: Stay green
SNP: Single-nucleotide polymorphism
Sp: Sub-populations
TKW: Thousand kernel weight
TPM: Transcripts per million Kb calculation

## Acknowledgments

The authors are grateful to Prof. R.A. McIntosh, Plant Breeding Institute, University of Sydney, for language editing and proofreading of the draft manuscript. We are very grateful to Dr Fengping Yuan of State Key Laboratory of Crop Stress Biology for Arid Areas, Mrs Haiying Wang of College of Horticulture, Northwest A&F University for assistance with DNA extraction and RNA extraction experiments.

## Funding

This study was supported by the National Key R&D Program of China (No. 2022YFE0116200), the Key R&D Program of Qinghai Province (2022-NK-125) and Key R&D Program of Yangling Seed Industry Innovation Center (grant no. Ylzy-xm-01).

## Author contributions

R.Y. designed the experiments; R.Y. and X.C. wrote the manuscript; R.Y., X.C., R.N., J.L., Y.H., and X.L. participated in the phenotypic measurements; J.W., Q.Z., C.Z., and M.Y. assisted in the data analysis and processing; J.W., Q.Z., W.Z., C.W., T.W., and B.S. revised the manuscript, while Z.K. and D.H. conceived and directed the project.

## Competing interests

The authors declare that they have no competing interests.

## Data Availability

The data presented in this study are available on reasonable request from the corresponding author.

## Supplementary files

**Table S1** Summary of the 565 worldwide wheat accessions.

**Table S2** The spectral indices used in this work.

**Table S3** Post-anthesis AT of three anthesis types of samples at different stages in two seasons.

**Table S4** Broad-sense heritability of SG traits.

**Table S5** Detailed information of all QTLs detected in each chromosome.

**Table S6** GO enrichment analysis of the 1,085 high-confidence candidate genes located in the QTL region.

**Table S7** SG variations distribution in worldwide wheat accessions about TraesCS6B03G0356400.

**Table S8** SG variations distribution in worldwide wheat accessions about TraesCS2B03G1299500.

**Table S9** SG variations distribution in worldwide wheat accessions about TraesCS2A03G1081100.

**Fig. S1** Daily air temperature and rainfall post-anthesis during the 2020-2021 (A) and 2021-2022 (B) growing seasons.

**Fig S2** Correlation coefficients between the different traits during the 2020-2021 (A) and 2021-2022 (B).

**Fig. S3** Relative stay green scores (RSGS) in 2021-2022 panel lines released pre- and post-1970. SG scores based on SG_NDVI_ and average SG scores in the panel at different release times are shown in lower left corner (A); SG score based on SG_GNDVI_ and average SG score in wheat panel at different release times are in the upper right corner (B); SG score based on SG_OSAVI_ and average SG score in wheat panel at different release times are shown in the lower left corner (C); SG score based on SG_NDRE_ and average SG score in wheat panel at different release times are shown in upper right corner (D).

**Fig S4** Manhattan plots of SG_NDVI1_, SG_NDVI2_, SG_NDVI3_ and SG_NDVI4_ in 2020-2021 (A-D, respectively), and 2021-2022 (E-H, respectively).

**Fig S5** Manhattan plots of SG_GNDVI1_, SG_GNDVI2_, SG_GNDVI3_ and SG_GNDVI4_ in 2020-2021 (A-D, respectively), and 2021-2022 (E-H, respectively).

**Fig S6** Manhattan plots of SG_NDRE1_, SG_NDRE2_, SG_NDRE3_ and SG_NDRE4_ in 2020-2021

(A-D, respectively), and 2021-2022 (E-H, respectively).

**Fig. S7** Manhattan plots of SG_OSAVI1_, SG_OSAVI2_, SG_OSAVI3_ and SG_OSAVI4_ in 2020-2021 (A-D, respectively), and 2021-2022 (E-H, respectively).

**Fig. S8** Frequency of QTL localization of multiple SI-based traits at multiple senescence stages. (S1 to S4, stage 1 to stage 4).

**Fig. S9** GO enrichment analysis of candidate genes in the SG QTL region.

**Fig. S10** Expression profile of *TraesCS6B03G0356400* (A), *TraesCS2B03G1299500* (B), and *TraesCS2A03G1081100* (C) at different time under heat stress 10 days after anthesis in the flag leaf. Shaannong 253 and Shannong 15 are SG material and non-SG material, respectively. CK means grown in the greenhouse with 22℃, 16 hours and 18℃, 8 hours; test means grown in the greenhouse with 38℃, 8hours and 24℃, 16hours.

